# PPAR-γ regulates the effector function of human T_H_9 cells by promoting glycolysis

**DOI:** 10.1101/2022.08.16.503972

**Authors:** Nicole L. Bertschi, Oliver Steck, Fabian Luther, Cecilia Bazzini, Leonhard von Meyenn, Andrea Felser, Irene Keller, Olivier Friedli, Stefan Freigang, Nadja Begré, Cristina Lamos, Max Philip Gabutti, Michael Benzaquen, Markus Laimer, Dagmar Simon, Jean-Marc Nuoffer, Christoph Schlapbach

## Abstract

T helper 9 cells (T_H_9) are key drivers of allergic tissue inflammation. They are characterized by the expression of type 2 cytokines, such as IL-9 and IL-13, and the peroxisome proliferator-activated receptor gamma (PPAR-γ) transcription factor. However, the functional role of PPAR-γ in human T_H_9 cells remains unknown. Here, we demonstrate that PPAR-γ drives activation-induced glycolysis, which, in turn, specifically promotes the expression of IL-9, but not IL-13, in an mTORC1-dependent manner. In vitro and ex vivo experiments on skin samples of allergic contact dermatitis showed that the PPAR-γ-mTORC1-IL-9 pathway was active in T_H_9 cells in human skin inflammation. Additionally, we found that tissue glucose levels were dynamically regulated in acute allergic skin inflammation, suggesting that in situ glucose availability is linked to distinct immunological signals in vivo. Furthermore, paracrine IL-9 induced the lactate transporter MCT1 in IL-9R^+^ T_H_ cells, where it increased aerobic glycolysis and proliferative capacity. Taken together, our findings delineate a hitherto unknown relationship between PPAR-γ-dependent glucose metabolism and the pathogenic effector function of human T_H_9 cells.

**Graphical abstract:** 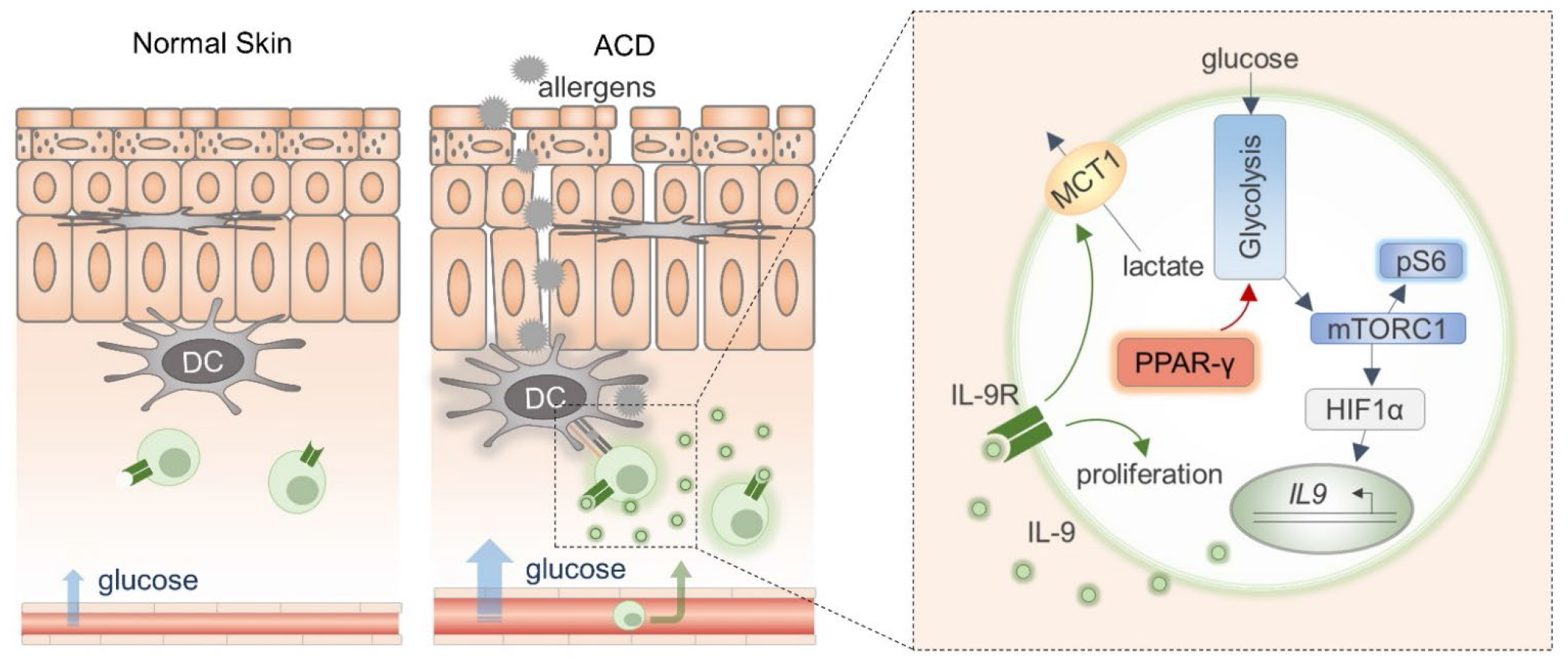

## Introduction

T helper (T_H_) cells have evolved into distinct subsets that mediate specific immune responses, protecting the host from various infectious and noninfectious challenges (*1*). However, impaired regulation of T_H_ cells can lead to inflammatory disease. Emerging evidence from both mice and humans indicates that type-2-driven diseases are mediated by a distinct subpopulation of the T_H_2 cells, referred to as pathogenic T_H_2 cells (pT_H_2 cells) (*2*), which, together with their effector molecules, serve as the prime targets for novel therapeutic approaches. In fact, pT_H_2 cells from a wide range of diseases such as allergic asthma, eosinophilic esophagitis (EoE), nasal polyps, and allergic contact dermatitis (ACD) share a common underlying transcriptome and overlapping functional characteristics (*3-9*). In particular, pT_H_2 cells express the peroxisome proliferator-activated receptor-γ (PPAR-γ) transcription factor and the IL-17RB and IL-9R cytokine receptors. They also secrete high levels of interleukin (IL-)13, IL-5, and IL-9 (*2*). Interestingly, PPAR-γ antagonism in human T_H_2 cells has been shown to inhibit the production of IL-9, while mice with T_H_ cell-specific PPAR-γ knockout are protected against T_H_2-mediated immunopathology (*10, 11*). This suggests that PPAR-γ plays an important functional role in pT_H_2 cells. Yet, despite the fact that PPAR-γ is intricately linked to the pT_H_2 cell phenotype, the mechanisms underlying the regulation of pT_H_2 cell function by PPAR-γ remain largely unknown.

The functional investigation of PPAR-γ in human T_H_ cells has been hampered by the low frequency of PPAR-γ-expressing T_H_ cells in human peripheral blood. However, we have recently identified IL-9-producing T_H_9 cells as a subpopulation of PPAR-γ^+^ T_H_2 cells that possesses the key characteristics of pT_H_2 cells (*8*). Human T_H_9 cells reside within the CCR4^+^/CCR8^+^ population of effector memory T cells (T_EM_). Functionally, they produce high levels of IL-9 and IL-13 and express transcription factors associated with the pT_H_2 lineage. Both in vitro and in vivo primed T_H_9 cells are distinguished from conventional T_H_2 (cT_H_2) cells: T_H_9 cells produce IL-9 in a transient activation-dependent manner and express PPAR-γ, which they rely on for their full effector function. Due to these key similarities shared with pT_H_2 cells, T_H_9 cells represent a valuable tool for studying the functional role of PPAR-γ in human T_H_ cell biology.

PPAR-γ is a ligand-activated nuclear receptor classically known for regulating lipid and glucose metabolism in adipocytes and other mesenchymal cells (*12*). PPAR-γ is activated by synthetic ligands, such as thiazolidinediones, as well as endogenous ligands thought to be derived from fatty acids (*13, 14*). Once activated, PPAR-γ dimerizes with the retinoid X receptor (RXR) and binds to genomic PPAR-responsive regulatory elements (PPREs) to control the expression of genes involved in lipid and glucose metabolism, as well as inflammation (*12*). The functional role of PPAR-γ in T_H_2 cells is best described in murine models, where it promotes IL-33R expression and thereby enhances the sensitivity of T_H_2 cells to tissue alarmins in allergic inflammation. Accordingly, mice with CD4-specific *Pparg* knockout exhibit impaired antiparasitic immunity, which protects them from allergic lung inflammation (*10, 11*). In humans, however, the function of PPAR-γ in pT_H_2 cells and its role in allergy are yet to be characterized.

Here, we sought to investigate the mechanism by which PPAR-γ regulates the effector function of human pT_H_2 cells, using in vitro and in vivo primed T_H_9 cells as an experimental tool. Collectively, our data point to the central role of PPAR-γ in promoting aerobic glycolysis, activating mTORC1, and stimulating IL-9 production in pT_H_2 cells and uncover a previously unknown functional role of paracrine IL-9 in promoting metabolic adaptation to high-glucose environments in acute allergic skin inflammation. Accordingly, PPAR-γ, IL-9, and their downstream targets might represent therapeutic leverage points in ACD and type-2-driven diseases.

## Results

### In vitro and in vivo primed T_H_9 cells display key features of pathogenic T_H_2 cells

The transcriptomic signature of pT_H_2 cells has been previously identified by single-cell analysis of T cells extracted from multiple T_H_2-driven diseases (*2*). To test whether human in vitro primed T_H_9 cells recapitulate the core pT_H_2 cell phenotype, we differentiated naïve T cells into T_H_1 (IL-12), T_H_2 (IL-4), T_H_9 (IL-4+TGF-β), and iT_REG_ (TGF-β) cells and performed transcriptomic profiling using RNA sequencing (RNA-seq) at day 7. Pairwise comparison to other subsets showed that 1492 genes were specifically upregulated in T_H_9 cells (Fig. 1A). We then compared our T_H_9 transcriptome with three pT_H_2-specific transcriptomes identified in EoE (*3*), allergic asthma (*4*), and allergen-specific T_H_2 cells, respectively (*6*) (Fig. 1B). *PPARG, IL5, IL17RB*, and *IL9R* were upregulated in all four datasets, while *IL9* was upregulated in three of the four. High levels of PPAR-γ were confirmed at the protein level in both in vitro and in vivo primed T_H_9 cells (Fig. 1C-F and Fig. S1A).

**Fig. 1:**
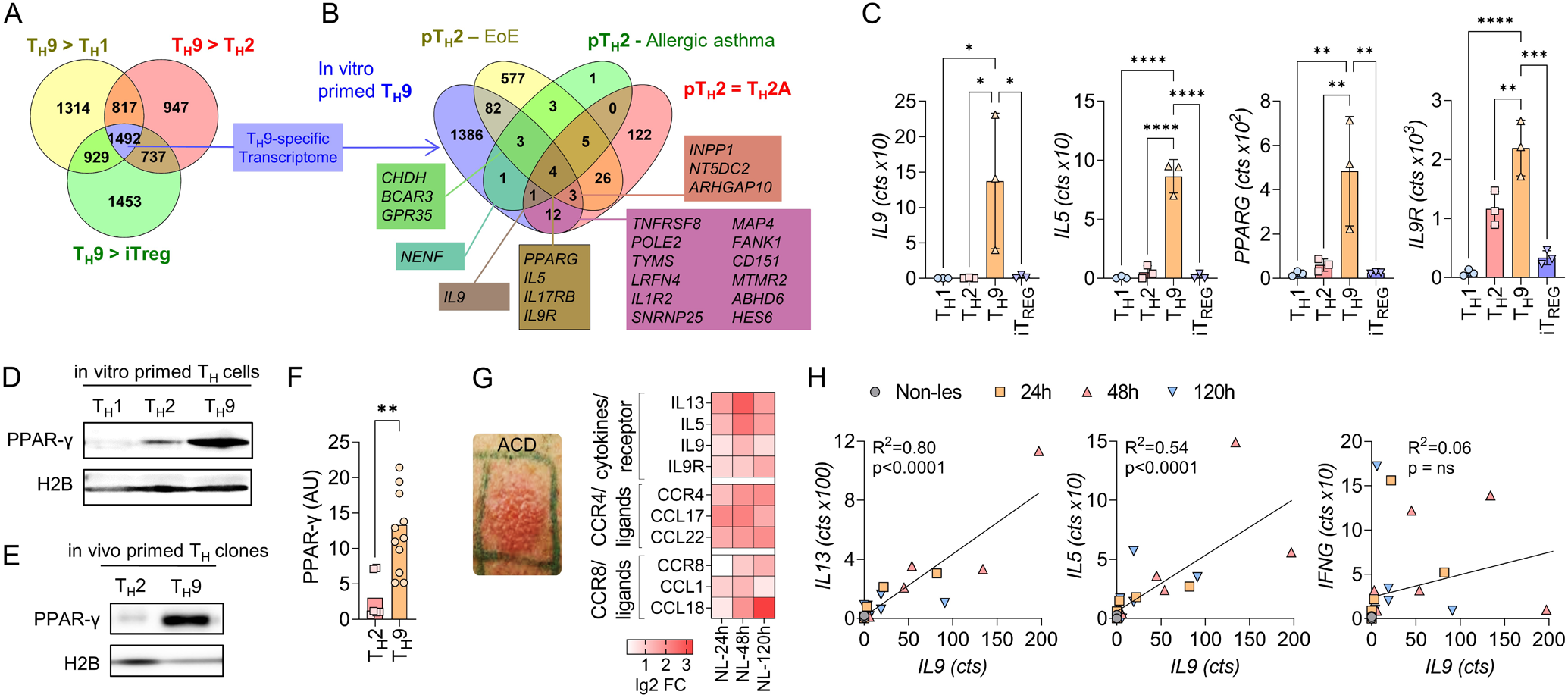
In vitro and in vivo primed T_H_9 cells display key features of pathogenic T_H_2 cells. **(A)** Venn diagram of RNA-seq data from T_H_ cell subsets primed in vitro showing the number of genes significantly upregulated in T_H_9 cells compared to other cell subsets (p_adj_ < 0.05). **(B)** Venn diagram of T_H_9-specific transcriptome identified in (A) and pT_H_2-associated genes identified in eosinophilic esophagitis (EoE) (*3*), allergic asthma (*4*), and allergen-specific T_H_2 cells (T_H_2A) (*6*). **(C)** Expression of selected pT_H_2-associated genes as determined in (A). **(D-F)** Western blot analysis of PPAR-γ in different T_H_ cell subsets primed in vitro (D) and in T_H_2 and T_H_9 clones primed in vivo (E, F). The data in (F) is expressed in arbitrary units (AU). **(G)** Changes in gene expression of selected pT_H_2-associated genes. **(H)** In-sample correlations of T cell cytokines with *IL9*. The data are representative of independent experiments with at least three (A-F) or six (G, H) donors. Statistics: (A) Differences between treatment groups were calculated as an adjusted log-fold change, and hypothesis testing was performed using the Benjamini-Hochberg adjusted p-value (DESeq2). (C) One-way ANOVA, followed by a Dunnett’s test for multiple comparisons. (F) Two-tailed unpaired t-test. (H) Simple linear regression. The data are presented as mean ± SD, *p<0.05, **p<0.01, ***p<0.001, ****p<0.0001.

As we had previously identified PPAR-γ^+^ T_H_9 cells in ACD (*8*), we next considered validating the association between the T_H_9 and pT_H_2 phenotypes in acute allergic skin inflammation. Time course transcriptomics of untreated non-lesional (NL) skin and positive patch test reactions of lesional skin to nickel at 24 h, 48 h, and 120 h post allergen application showed a marked upregulation of the pT_H_2-expressed genes in ACD (Fig. 1G). Across individual samples, the expression of *IL9* correlated with the expression of *IL13* (R^2^ = 0.80; P < 0.0001), *IL5* (R^2^ = 0.54; P < 0.0001), *IL31* and *IL19*, but not *IFNG* (Fig. 1H and Fig. S1B), consistent with the predominance of T_H_9 cells in the T_H_2 cell pool. Due to the fact that *PPARG* is expressed in various skin cell types, including keratinocytes (*8*), the correlation analysis of *PPARG* with *IL9* does not allow any conclusion with regard to T_H_9 cells.

Collectively, these data show that both in vitro and in vivo primed T_H_9 cells express the core features of pT_H_2 cells, including upregulated expression of PPAR-γ, IL-5, IL-9, and IL-9R. They can thus serve as model cells for studying the functional role of PPAR-γ in human T_H_ cells. Further, human ACD appears to be a valid model for studying the functionality of T_H_9 cells ex vivo.

### PPAR-γ mediates the high glycolytic activity of T_H_9 cells

To investigate the role of PPAR-γ in human T_H_ cells, we first assessed the transcriptional response of activated T_H_9 clones to treatment with GW9662, a potent PPAR-γ antagonist. Pathway analysis of RNA-seq data revealed concerted downregulation of genes associated with T cell activation, glucose metabolism, and aerobic glycolysis (Fig. S2A). At the single gene level, mRNA expression of all aerobic glycolysis enzymes was significantly downregulated by PPAR-γ inhibition (Fig. 2, A and B). This prompted us to further analyze the role of PPAR-γ in aerobic glycolysis of T_H_9 cells. In vitro primed T_H_9 cells showed higher glycolytic activity than T_H_1- or T_H_2-primed T cells after activation with αCD3/CD2/CD28 (Fig. 2C). To verify whether PPAR-γ was involved in this process, we starved in vitro primed T_H_9 cells in glucose-free medium in presence or absence of GW9662. We then performed real-time ECAR measurements before and after activation with αCD3/CD2/CD28 in either low or high-glucose environments. PPAR-γ inhibition hampered the glycolytic response in high-but not low-glucose environments, particularly following T cell activation (Fig. 2, D and E). These findings were corroborated by measurements of glucose uptake, in which in vitro primed T_H_9 cells showed higher glucose uptake compared to T_H_0, T_H_1, and T_H_2 cells (Fig. 2F). Importantly, glucose uptake in T_H_9 cells was PPAR-γ-dependent (Fig. 2G). The findings above were further confirmed using an alternative PPAR-γ antagonist, T0070907 (Fig. S2B).

**Fig. 2:**
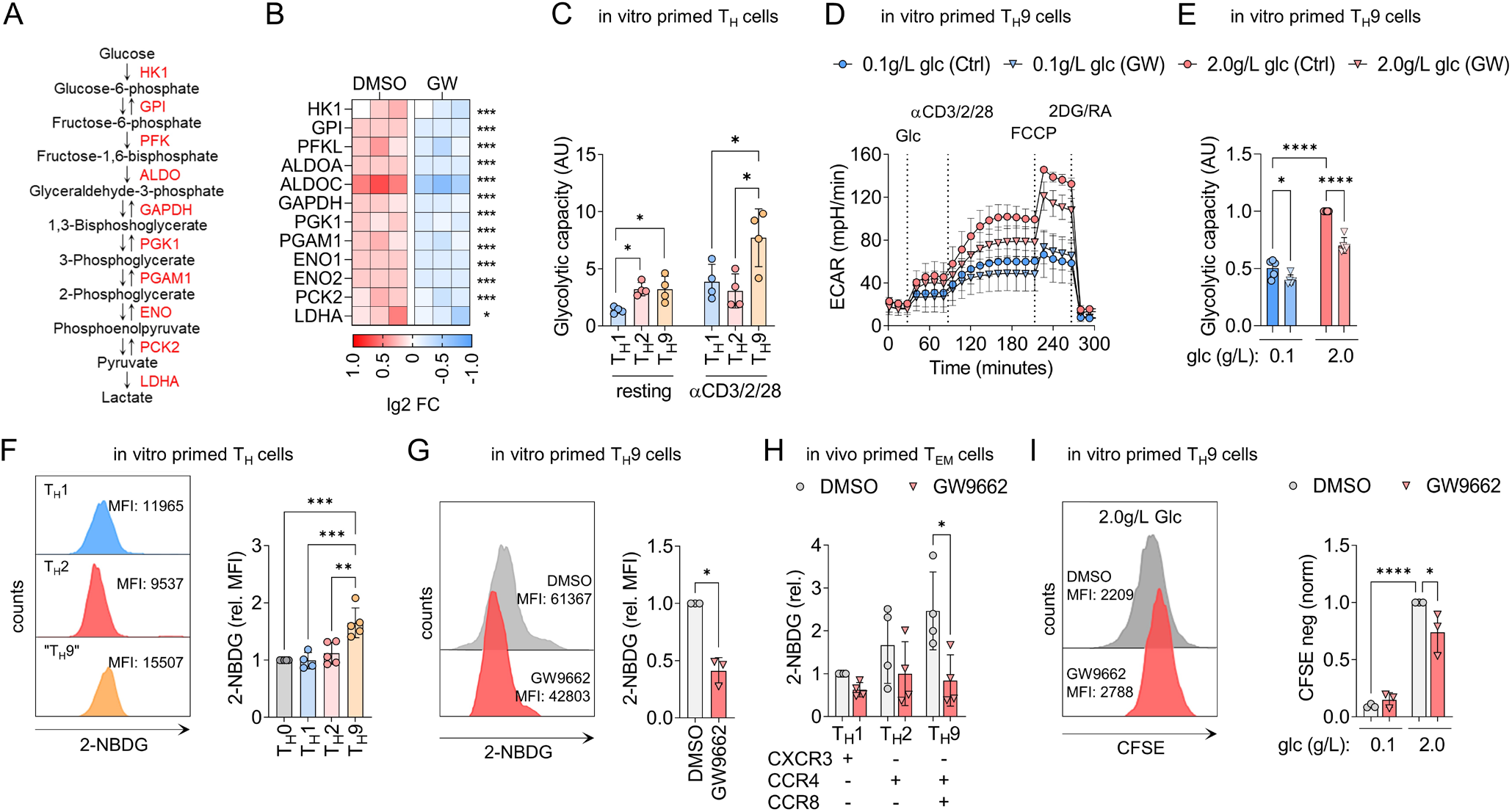
PPAR-γ mediates the high glycolytic activity of T_H_9 cell. **(A)** Intermediates and enzymes (red) of aerobic glycolysis. **(B)** RNA-seq of T_H_9 clones incubated in presence of GW9662 for 48 h and activated by αCD3/CD2/CD28 for 12 h. **(C)** Maximal glycolytic capacity of in vitro primed T_H_ cells in the resting state (left) and 24 h after activation with αCD3/CD2/CD28 (right). **(D)** ECAR measurements of in vitro primed T_H_9 cells cultured in media with glucose of different levels and GW9662 for 48 h and activated by injection of glucose and αCD3/CD2/CD28. **(E)** Glycolytic capacity of in vitro primed T_H_9 cells from (D). **(F)** Glucose uptake by in vitro primed T_H_ cells measured with fluorescent 2-NBDG uptake by flow cytometry. **(G)** Glucose uptake by naïve T_H_ cells primed under T_H_9 conditions for 7 days in the presence or absence of GW9662. **(H)** Glucose uptake of in vivo primed effector memory T_H_ cells (T_EM_) (CD45RA^-^/CCR7^-^/CD25^-^) sorted by flow cytometry into T_H_1, T_H_2, and T_H_9 cells according to their chemokine receptor profile. Sorted T_H_ cells were incubated in presence or absence of GW9662 for 48 h, and activated by αCD3/CD2/CD28 for 4 h. **(I)** Proliferation of in vitro primed T_H_9 cells, activated by αCD3/CD2/CD28 for 4 days in the presence or absence of GW9662 and glucose of different levels, measured with CFSE dilution by flow cytometry. The data are representative of one experiment with three clones from one donor (B) or independent experiments with two (F), three (D, E, G, I), or four donors (C, H). Statistics: (B) Differences between treatment groups were calculated as an adjusted log-fold change, and hypothesis testing was performed using the Benjamini-Hochberg adjusted p-value (DESeq2). (C, F, H) One-way ANOVA, followed by a Tukey’s test for multiple comparisons. (G) Two-tailed paired t-test. (E, I) One-way ANOVA, followed by a Šidák’s test for multiple comparisons. The data are presented as mean ± SD, *p<0.05, **p<0.01, ***p<0.001, ****p<0.0001.

We next expanded our findings to in vivo primed memory T_H_ cells, leveraging the ability to sort different subsets ex vivo based on their chemokine receptor profile, including CXCR3^−^/CCR4^+^/CCR6^−^/CCR8^+^ T_H_9 cells (*8, 15*). Consistent with our previous observations, in vivo primed T_H_9 cells showed higher glucose uptake compared to other T_H_ cell subsets, and GW9662 significantly inhibited the glycolytic activity in T_H_9 cells but not in T_H_2 or T_H_1 cells (Fig. 2H).

Given the central role of aerobic glycolysis for T cell proliferation (*16*), we finally tested whether PPAR-γ inhibition affected TCR stimulation-induced proliferation in low and high-glucose environments. Both GW9662 and T0070907 significantly reduced αCD3/CD2/CD28-induced proliferation in high-glucose environments (Fig. 2I and Fig. S2C).

Collectively, these data strongly suggest that both in vitro and in vivo primed human T_H_9 cells are characterized by high glycolytic capacity post-TCR-stimulation. Moreover, PPAR-γ signaling appears to be a crucial mediator of this process.

### High glycolytic activity in T_H_9 cells differentially regulates cytokine expression

PPAR-γ is required for the full effector function in T_H_9 and pT_H_2 cells, including the production of proinflammatory cytokines (*8, 10, 11*). Based on our findings, we hypothesized that PPAR-γ might control cytokine production indirectly, namely by promoting glycolysis. To test this, we cultured in vitro primed T_H_9 cells in media containing different glucose concentrations and measured their cytokine profiles after activation. Production of the pT_H_2 marker cytokine IL-9 showed a strong dependency on glucose availability, whereas production of IL-13, the conventional T_H_2 cytokine, did not (Fig. 3A). Further, in vivo primed T_H_9 clones cultured in different glucose concentrations downregulated the production of IL-9 but not IL-13 in low-glucose environments (Fig. 3B).

**Fig. 3:**
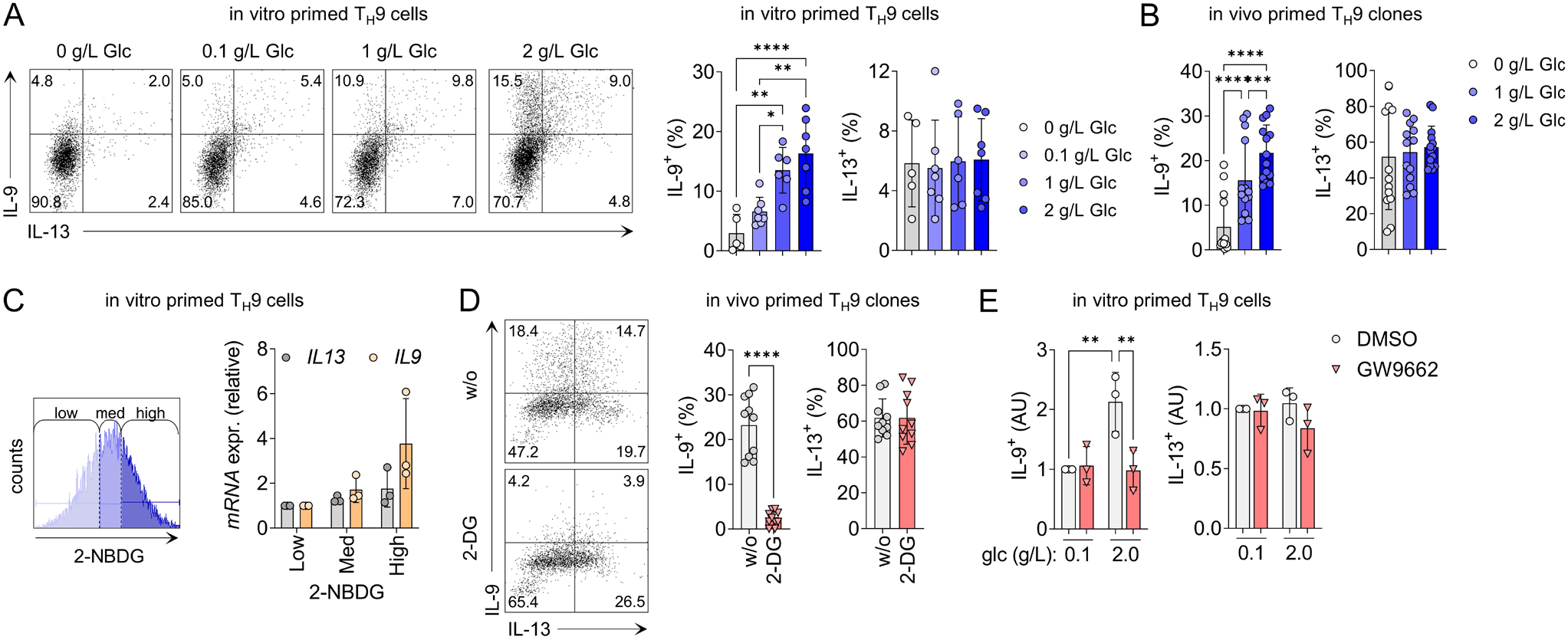
High glycolytic activity of T_H_9 cells regulates specific effector functions. **(A)** Cytokine expression of in vitro primed T_H_9 cells cultured for 48 h in media containing glucose of different levels, measured by flow cytometry 18 h after activation with αCD3/CD2/CD28. **(B)** Cytokine expression of in vivo primed T_H_9 clones cultured for 72 h in media containing glucose of different levels, measured as in (A). **(C)** In vitro primed T_H_9 cells were activated for 4 h with αCD3/CD2/CD28, then sorted by flow cytometry based on their glucose uptake measured by 2-NBDG uptake (left). Cytokine expression in the sorted T_H_ cell populations was measured by RT-qPCR (right). **(D)** Cytokine expression of in vivo primed T_H_9 cells cultured for 7 days in the presence of 2-DG, measured as in (A). **(E)** Cytokine expression of in vitro primed T_H_9 cells cultured for 48 h in media containing glucose of different levels and in the presence or absence of GW9662, measured as in (A). The data are representative of independent experiments with three (C, D, E) or six (A, B) donors. Statistics: (A) One-way ANOVA, followed by a Tukey’s test for multiple comparisons. (B, E) One-way ANOVA, followed by a Šidák’s test for multiple comparisons (D) Two-tailed paired t-test. The data are presented as mean ± SD, *p<0.05, **p<0.01, ***p<0.001, ****p<0.0001.

To demonstrate a direct relationship between high glucose metabolism and cytokine production, we next sorted in vitro primed T_H_9 cells according to their glucose uptake level, measured by the uptake of 2-NBDG. Subsequently, we performed RT-qPCR for *IL9* and *IL13*. Glucose uptake correlated with the expression of *IL9* but not *IL13* (Fig. 3C). We then inhibited glycolysis in T_H_9 cells using the glucose analog 2-deoxy-d-glucose (2-DG). In T_H_9 cells primed in vivo, 2-DG completely inhibited the expression of IL-9 but not IL-13 (Fig. 3D). Finally, PPAR-γ antagonism in high-glucose environments reduced the production of IL-9 to the levels seen in low-glucose environments, whereas IL-13 levels remained unaffected neither by PPAR-γ antagonism nor by low glucose availability (Fig. 3E). Taken together, these observations indicate a dichotomous role of glycolytic activity in regulating the production of IL-9 and IL-13 by activated T_H_9 cells.

### mTORC1 integrates glycolytic activity with the effector function in T_H_9 cells

We next investigated the mechanisms underlying the association between glycolysis and cytokine production in activated T_H_9 cells. Mammalian target of rapamycin complex 1 (mTORC1) is a central regulator of cellular metabolism and effector functions in T cells. Nutrients, such as glucose, are critical activators of mTORC1 (*17*). Furthermore, the mTORC1-hypoxia-inducible factor-1α (HIF-1α) pathway is necessary for the expression of IL-9 in murine T cells, with HIF-1α binding directly to the *Il9* promoter and activating its transcription (*18-20*). Given the role of the established mTORC1-HIF-1α-IL-9 axis and our previous results, we hypothesized that mTORC1 might mediate the PPAR-γ-dependent expression of IL-9 in T_H_9 cells.

Phosphorylation of mTORC1 in activated T_H_9 cells measured by phosphorylated S6 (pS6) was glucose-dependent and reduced by PPAR-γ inhibition under high glucose conditions (Fig. 4A, and Fig. S4, A, and B). Indeed, IL-9^+^ T cells were strongly enriched in the pS6^+^ cell population, whereas the proportion of IL-13^+^ T cells was similar in pS6^−^ and pS6^+^ populations (Fig. 4, B and C). Moreover, rapamycin-induced inhibition of mTORC1 decreased the production of IL-9 but not IL-13 in activated T_H_9 cells (Fig. 4, D and E, and Fig. S4C).

**Fig. 4:**
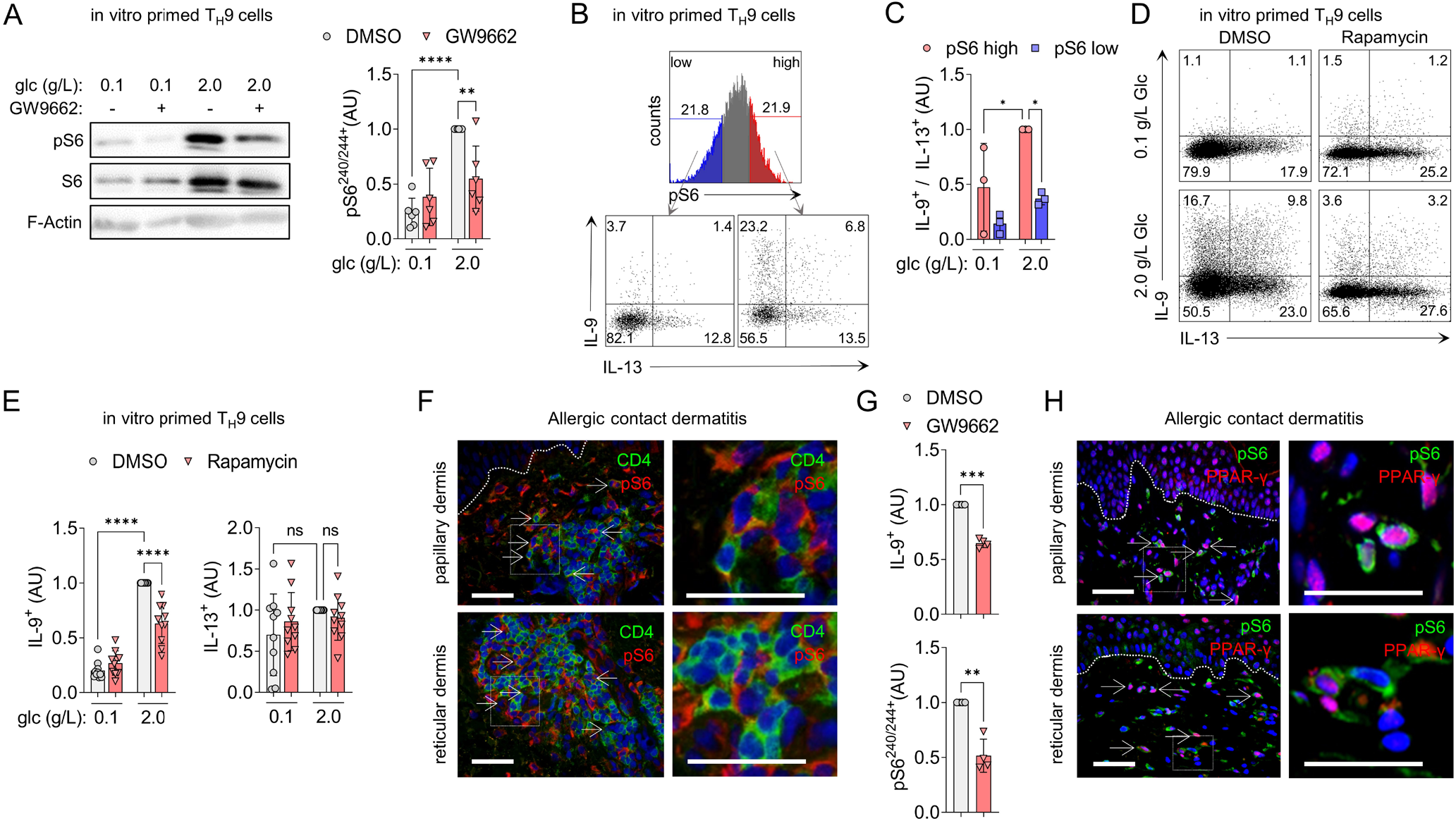
mTORC1 integrates bioenergetics with the effector function in T_H_9 cells. **(A)** Western blot analysis of pS6 and total S6 in T_H_9 cells primed in vitro. **(B)** In vitro primed T_H_9 cells were cultured in glucose of different levels for 48 h, and pS6 and IL-9 were measured 18 h after activation with αCD3/CD2/CD28 by flow cytometry. The histogram shows pS6 positive cells, split into high and low pS6. Dot-plots represent IL-9^+^ / IL-13^+^ clusters. **(C)** The IL-9^+^ / IL-13^+^ ratio in T_H_9 cells primed in vitro from (B). (**D-E)** Cytokine expression of in vitro primed T_H_9 cells after incubation in glucose of different levels and rapamycin for 48 h, measured as in (B). **(F)** Immunofluorescence staining for CD4 and pS6 on skin samples of allergic contact dermatitis (ACD). **(G)** Cytokine expression of T cells isolated from ACD skin biopsies incubated with GW9662 for 48 h, measured as in (B). **(H)** Immunofluorescence staining for pS6 and PPAR-γ on skin samples of ACD. The data are representative of independent experiments with at least one (F, G, H), three (C), six (A) donors or nine (E) donors. Statistics: (A, C, E) One-way ANOVA, followed by a Šidák’s test for multiple comparisons. (G) Two-tailed paired t-test. The data are presented as mean ± SD, *p<0.05, **p<0.01, ***p<0.001, ****p<0.0001.

To verify whether T_H_9 cells expressed activated mTORC1 in human skin inflammation, we performed immunofluorescence staining on skin samples of ACD and isolated T cells from such lesions. Double immunofluorescence revealed that CD3^+^ and CD4^+^ T_H_ cells expressed pS6 in the infiltrate of ACD (Fig. 4F and Fig. S4D). T cells isolated from ACD incubated with GW9662 ex vivo showed a reduced activation of mTORC1 and a significantly inhibited IL-9 production (Fig. 4G). As it is known that ACD skin is infiltrated by a substantial number of PPAR-γ^+^ T_H_ cells (*8*), we finally investigated whether those cells would show the activation of mTORC1. Indeed, PPAR-γ^+^pS6^+^ double-positive lymphocytes were readily found throughout the dermis of ACD (Fig. 4H).

Collectively, these findings strongly suggest that glucose- and PPAR-γ-dependent production of IL-9 in T_H_9 cells is regulated via mTORC1 in acute allergic skin inflammation.

### Paracrine IL-9 promotes aerobic glycolysis in IL-9R^+^ T_H_ cells by inducing the lactate transporter MCT1

After revealing the association between PPAR-γ-dependent glycolytic activity and IL-9 production in T_H_9 cells, we hypothesized that paracrine IL-9 might regulate glucose metabolism and downstream effector functions in IL-9R^+^ T_H_ cells. Previous data suggested that *IL9R* is preferentially expressed in pT_H_2 and T_H_9 cells (Fig. 1 and refs. (*3, 4, 6, 7*)), but these findings had to be confirmed at the protein level and in human skin inflammation. Thus, we first confirmed the expression of IL-9R in T_H_9 clones primed in vivo (Fig. 5A), circulating CXCR3^−^/CCR4^+^/CCR8^+^ effector memory T_H_ cells (Fig. 5B), and T_H_ cells infiltrating lesions of ACD (Fig. 5, C and D), showing that pT_H_2 and T_H_9 cells are important targets of IL-9 in human skin inflammation. Next, we performed RNA-seq of IL-9R^+^ T_H_ clones and IL-9R^+^ T_H_ cells isolated from ACD skin biopsies, incubated with or without IL-9. The pathway analysis of IL-9-induced genes showed a coordinated induction of genes involved in aerobic glycolysis (Fig. 5E), most prominently *SLC16A1* (Fig. 5F). *SLC16A1* encodes the monocarboxylate transporter 1 (MCT1), enabling the rapid export of lactate across the plasma membrane and thereby exerting a glycolytic flux-controlling function (*21*). IL-9-induced expression of MCT1 in T_H_9 clones was confirmed at the protein level (Fig. 5G). Next, we investigated the expression and regulation of *SLC16A1* in different T_H_ cell populations. In vitro T_H_9 differentiation induced higher levels of *SLC16A1* than T_H_1, T_H_2, or iT_REG_ differentiation (Fig. 5H). Time course transcriptomics of TCR-stimulated T_H_ clones (*8*) revealed a significantly higher expression of *SLC16A1* in T_H_9 clones than in T_H_1 and T_H_2 clones (Fig. 5I), as well as a close correlation between the expression of *IL9* and *SLC16A1* (Fig. 5J). Finally, a Seahorse experiment confirmed that IL-9R^+^ T_H_ clones stimulated by IL-9 exhibited strongly elevated ECAR, in line with IL-9-dependent induction of active aerobic glycolysis and efficient cellular export of lactate (Fig. 5K). Accordingly, IL-9 increased extracellular lactate levels in cultured IL-9R^+^ T_H_ clones, and these levels were suppressed by the addition of BAY-8002, a potent MCT1 antagonist (MCT1-i) (Fig. 5L).

**Fig. 5:**
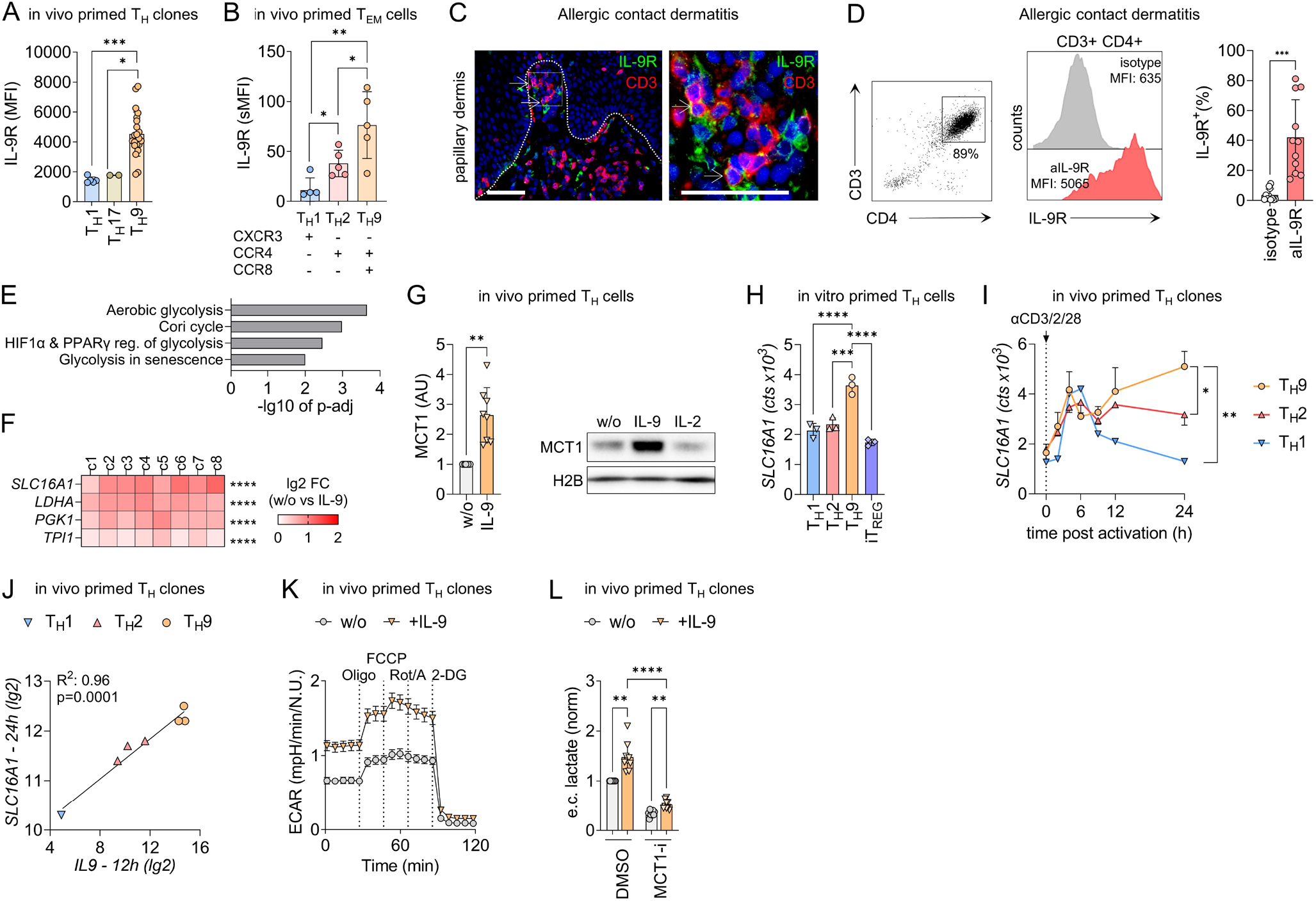
Paracrine IL-9 promotes aerobic glycolysis in IL-9R^+^ T_H_ cells by inducing the lactate transporter SLC16A1. **(A)** IL-9R levels in T_H_1, T_H_17, and T_H_9 clones primed in vivo using flow cytometry. **(B)** IL-9R levels were analyzed by flow cytometry in Peripheral Blood Mononuclear Cells (PBMCs) isolated from blood and stained for their chemokine receptor profiles. **(C)** Immunofluorescence staining for CD3 and IL-9R on skin samples of allergic contact dermatitis (ACD). **(D)** IL-9R levels of T cells isolated from ACD skin biopsies were determined by flow cytometry. Gating of CD3^+^CD4^+^ cells (left), representative histogram of the antibody and isotype (middle), and aggregated data (right). **(E)** Pathway analysis of the 250 most significant IL-9-induced genes from RNA-seq data of IL-9R^+^ T_H_ cells, isolated from blood and ACD skin biopsies, incubated with IL-9 for 12 h. **(F)** Changes in the expression of selected aerobic glycolysis genes in presence of IL-9 in IL-9R^+^ T_H_ cells from (E). **(G)** Western blot analysis of MCT1 expression in IL-9R^+^ T_H_ clones incubated with IL-9 and IL-2 for 48 h. **(H)** The RNA expression level of *SLC16A1* in T_H_ cells primed in vitro under T_H_1, T_H_2, T_H_9, or iT_REG_ priming conditions for 7 days. **(I-J)** Time course transcriptomic data of TCR-stimulated T_H_ clones (*8*) shows RNA expression levels of *SLC16A1* in (I) and correlation between the expression of *IL9* and *SLC16A1* in (J). **(K)** In vivo primed IL-9R^+^ T_H_ clones incubated with IL-9 for 16 h and analyzed by extracellular flux measurements. **(L)** IL-9R^+^ T_H_ clones incubated for 48 h with IL-9 and MCT1 inhibitor (MCT1-i) BAY-8002, respectively. Extracellular (e.c.) lactate levels were determined with the Lactate-Glo^™^ Assay (Promega). The data are representative of one experiment with one (C, I, K), two (A, T_H_17), four (A, T_H_1), or twenty-two (A, T_H_9) clones from one donor or independent experiments with at least three clones from one, (G), two (E, F, L) or three donors (G, H) or independent experiments with five (B) or eleven (D) donors. Statistics: (A) Two-tailed unpaired t-test. (B, D, G) Two-tailed paired t-test. (F) Differences between treatment groups were calculated as an adjusted log-fold change, and hypothesis testing was performed using the Benjamini-Hochberg adjusted p-value (DESeq2). (H) One-way ANOVA, followed by a Dunnett’s test for multiple comparisons. (J) Simple linear regression. (I, L) One-way ANOVA, followed by a Tukey’s test for multiple comparisons. The data are presented as mean ± SD, *p<0.05, **p<0.01, ***p<0.001, ****p<0.0001.

Taken together, these data indicate that paracrine IL-9 promotes aerobic glycolysis in IL-9R^+^ T_H_ cells by inducing the lactate transporter MCT1, which controls glycolytic flux.

### IL-9 promotes T cell proliferation in high-glucose environments

Considering the crucial role of aerobic glycolysis in supporting T cell proliferation (*22*), we next investigated the functional effects of IL-9-induced MCT1 expression on T cell proliferation. IL-9 induced strong proliferative responses in IL-9R^+^ T_H_ clones (Fig. 6A), and this proliferative boost was reversed by adding BAY-8002, the MCT1 inhibitor (Fig. 6B). Moreover, IL-9-induced proliferation was dependent on available glucose levels (Fig. 6C), which further supports the notion that IL-9 promotes glycolytic flux in IL-9R^+^ T_H_ cells to facilitate proliferation.

**Fig. 6:**
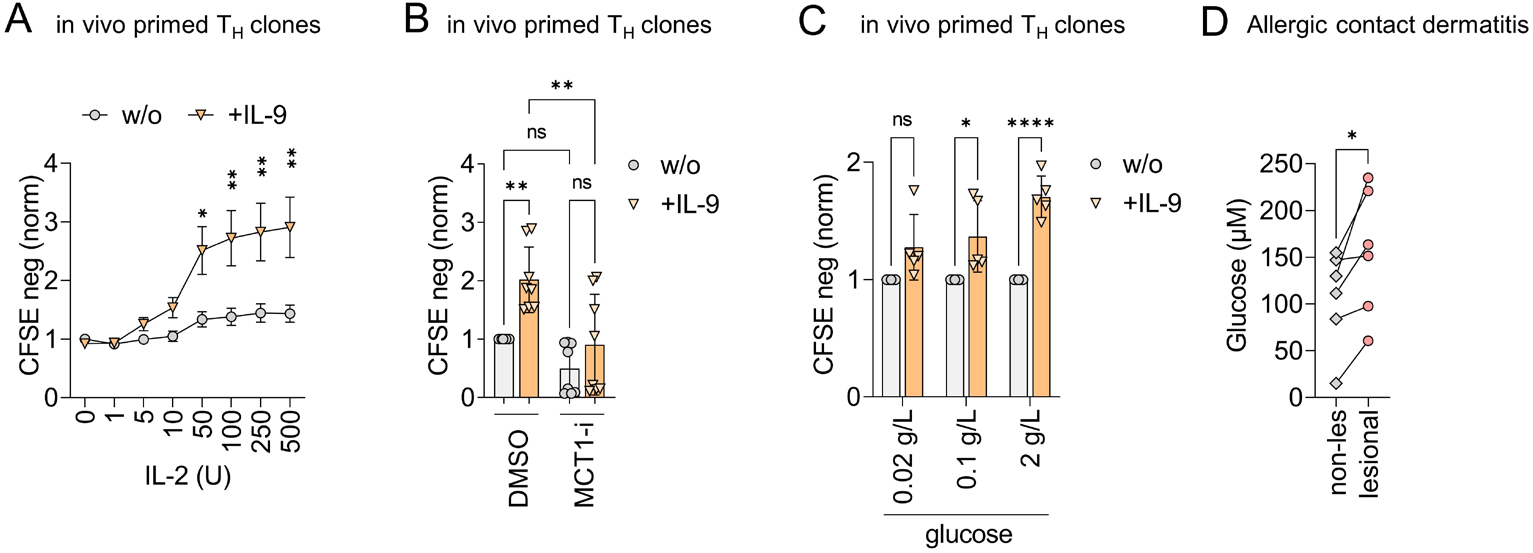
IL-9 promotes T cell proliferation in the high-glucose environment of allergic contact dermatitis. **(A)** Proliferation of in vivo primed IL-9R^+^ T_H_ clones in presence of IL-9 and different concentrations of IL-2, measured with CFSE dilution by flow cytometry after 3 days. **(B)** Proliferation of in vivo primed IL-9R^+^ T_H_ clones in presence of IL-9 and MCT1 inhibitor (MCT1-i) BAY8002 measured as in (A). **(C)** In vivo primed IL-9R^+^ T_H_ clones were cultured in media containing glucose of different levels for 7 days. Proliferation was measured in presence of IL-9 with CFSE dilution by flow cytometry as in (A). **(D)** Glucose concentrations measured with the Glucose-Glo^™^ Assay (Promega) of interstitial fluids of lesional skin of positive patch test reactions to different allergens (Supplementary Table S3) and adjacent non-lesional skin biopsies. The data are representative of one experiment with four clones from one donor (A, C) or independent experiments with eight clones from one donor (B) or independent experiments with six donors (D). Statistics: (A, C) Two-way ANOVA, followed by a Šidák’s test for multiple comparisons. (B) One-way ANOVA, followed by a Tukey’s test for multiple comparisons. (D) Two-tailed paired t-test. The data are presented as mean ± SD, *p<0.05, **p<0.01, ***p<0.001, ****p<0.0001.

Finally, we investigated whether tissue glucose levels are dynamically regulated in acute allergic skin inflammation, in which T_H_9 cells have been shown to highly express IL-9 (*8, 23*). To this end, we determined glucose levels in the interstitial fluid of tissue homogenates from non-lesional and lesional ACD skin, respectively. The interstitial fluid of the lesional skin contained higher glucose levels than the matched non-lesional skin samples (Fig. 6D), which suggests that IL-9-related human allergic skin inflammation is associated with changes in glucose availability in vivo.

Collectively, these observations suggest that paracrine IL-9 facilitates the proliferation of T cells by stimulating aerobic glycolysis through the induction of the lactate transporter MCT1, possibly contributing to the proliferation of IL-9R^+^ T_H_ cells in the high-glucose environment of ACD and acute allergic tissue inflammation.

## Discussion

Here, we used in vivo and in vitro primed T_H_9 cells as a model PPAR-γ^+^ T cell population to study the role of PPAR-γ in human T_H_ cells, particularly in disease-associated pT_H_2 cells. We provide evidence that PPAR-γ is a positive regulator of aerobic glycolysis in activated human T_H_9 cells. High glycolytic activity, in turn, was found to differentially regulate the effector function, particularly the expression of IL-9. Paracrine IL-9 signaling enhanced the expression of the lactate transporter MCT1, which provided a proliferative advantage to IL-9R^+^ T_H_ cells in high-glucose environments. Our data point to a previously unknown role of PPAR-γ and IL-9 in facilitating immunometabolic sensing of the tissue microenvironment. In the future, the PPAR-γ-mTORC1-IL-9 axis can be evaluated as a therapeutic target for type-2-driven skin inflammation and pT_H_2-mediated disease.

*PPARG* is highly and specifically expressed in pT_H_2 cells isolated from a variety of human type-2-driven diseases (*3-9*). In mice, PPAR-γ is functionally important for the pathogenicity of T_H_2 cells in allergic tissue inflammation, where it regulates the expression of IL-33R and increases the sensitivity of T_H_2 cells to the tissue alarmin IL-33 (*10, 11*). While IL-33R (*IL1RL1*) is also associated with the pT_H_2 phenotype in humans (*3, 5-7*), we did not find that PPAR-γ regulates its expression in T_H_9 cells, nor did we observe the reactivity of T_H_9 cells to IL-33 (data not shown). However, we found that PPAR-γ promotes the expression of glycolytic enzymes in activated T_H_9 cells, thereby supporting the crucial metabolic switch to aerobic glycolysis that early T cell activation and proliferation depend on (*24, 25*). Our findings align with a recent report of genome-wide CRISPR screens in T_H_ cells, where PPARG, together with GATA3, IRF4 and BATF, was found to form a core T_H_2 regulatory network (*26*). In that analysis, PPAR-γ was identified as particularly important for the activation of T_H_2 cells but to a lesser extent for their differentiation. This may be related to the role of PPAR-γ in supporting aerobic glycolysis in pT_H_2 cells, as observed in the present study. In fact, both rapid upregulation of aerobic glycolysis (*18*) and sustained cell proliferation after activation (*27*) have been previously described in murine T_H_9 cells. Our data now suggest that this is, at least in part, a PPAR-γ-dependent process, providing a functional explanation for the association of PPAR-γ with the pT_H_2 phenotype. Functionally, PPAR-γ might thus provide pT_H_2 cells with an advantage over neighboring cells in competition for critical nutrients, such as glucose, in acute allergic tissue inflammation.

Immunometabolism is increasingly appreciated as an essential regulator of the T cell effector function (*22*). However, the full spectrum of effector functions regulated by glycolytic activity in T_H_ cells remains incompletely understood (*28*). Previously, we found a differential effect of PPAR-γ antagonism on T_H_2 cytokine profiles, with IL-9 expression dependent on PPAR-γ-signaling while IL-13 being expressed independently thereof (*8*). In the present study, we identified glycolytic activity and sequential mTORC1 activation as a mechanistic link between PPAR-γ and IL-9 expression. Indeed, mTORC1 has been shown to promote IL-9 expression in murine T_H_ cells by activating HIF-1α, which has a direct binding site on the *IL9* promoter (*18*). Therefore, our study further supports the notion that distinct metabolic pathways can have a selective effect on the T_H_ cell effector function and, in principle, demonstrates that these immunometabolic links could be leveraged for targeted manipulation of immune function (*29*).

Our finding of increased interstitial glucose levels in acute skin inflammation is of particular interest, as the skin is generally considered a low-glucose environment (*30*). A recent study in mice has also found increased tissue glucose levels in type-2-mediated lung inflammation and has linked increased glycolytic activity to pathogenic functions of type 2 innate lymphoid cells (ILC2) (*31*). Thus, pT_H_2 cells might be particularly adapted to function in the high-glucose environment of acute allergic tissue inflammation due to the high glycolytic capacity provided by PPAR-γ.

The specific dependence of IL-9 production on active glycolysis in T_H_9 cells raises the question of why a defined immunological signal depends on the availability of a particular nutrient. The putative answer comes from our finding that IL-9 induces MCT1 expression, enabling efficient lactate export and NAD^+^ regeneration for further glycolytic activity via GAPDH. Indeed, inhibition of MCT1 during T cell activation selectively inhibits early T cell proliferation (*32*) and has been proposed as a novel approach for immunosuppressive therapy (*33*). IL-9 is a pleiotropic proinflammatory cytokine in allergic inflammation, but its target cells and mechanism of action remain not fully defined (*34-36*). Our data now suggest a previously unrecognized role of IL-9 in inducing lactate transport capacity in IL-9R^+^ T_H_ cells, whereby it increases their glycolytic capacity and proliferative potential. Interestingly, *SLC16A1* expression is also upregulated in lesional skin of atopic dermatitis and forms part of a dynamic immune signature that reflects progressive immune activation dominated by T_H_2 cells (*37*).

Our study has several limitations and raises intriguing questions that remain to be addressed. For example, it remains unknown to what extent the glucose concentrations used in our in vitro assays are representative of the in vivo metabolic environment. Experimental modulation of interstitial levels of ubiquitous metabolites, such as glucose, remains challenging even in animal models (*38*). The most critical metabolic pathways for T cell function have been identified in reductionist in vitro assays. Furthermore, how pT_H_2 cells compete with neighboring tissue cells for nutrients, such as glucose, remains to be addressed. It is worth investigating whether PPAR-γ indeed confers a competitive metabolic advantage to pT_H_2 cells in vivo and whether such differences in metabolic fitness translate into functional outcomes that can be targeted therapeutically. Finally, it remains to be investigated whether the PPAR-γ-mTORC1-IL-9 axis and its downstream target MCT1 are viable therapeutic targets for allergic skin inflammation. Collectively, our findings encourage further research into the molecular details of how PPAR-γ regulates the metabolism and function of pT_H_2 cells to enable the development of novel therapeutic interventions.

## Materials & Methods

### Study and experimental design

The aim of this study was to investigate the mechanism by which PPAR-γ regulates the effector function of human pT_H_2 cells. To this end, we used in vitro and in vivo primed T_H_9 cells as a model of PPAR-γ^+^ T cell population and we performed RNA-seq analysis of activated T_H_9 clones upon treatment with the PPAR-γ antagonist GW9662. This analysis pointed to a role of PPAR-γ in the regulation of glycolytic activity in T_H_9 cells. This hypothesis was tested in vitro by measuring real time extracellular acidification rate using Seahorse Analyzer as well as glucose uptake, proliferation and cytokines expression by flow cytometry, in low and high-glucose environments and/or in presence or absence of the PPAR-γ inhibitor. We next investigated the ability of mTORC1 to sense glucose availability and mediate the PPAR-γ-dependent IL-9 expression by assessing S6 phosphorylation on cell-based experiments but also on skin samples of ACD, on which immunofluorescence staining was performed. Further, we tested whether IL-9 regulates glucose metabolism and downstream effector functions in IL-9R^+^ T_H_ cells by performing RNA-seq in presence or absence of IL-9. The analysis revealed that IL-9 stimulated T_H_ cells showed an induction of genes involved in aerobic glycolysis, which was confirmed at the protein level. The effect of IL-9 in increasing glycolytic capacity and proliferative potential of IL-9R+ T_H_ cells was analyzed by Seahorse experiments and CFSE dilution using flow cytometry, respectively. Finally, to evaluate how tissue glucose levels are dynamically regulated in acute allergic skin inflammation, interstitial fluid of tissue homogenates from non-lesional and lesional skin of ACD were analyzed with a glucose detection luminescence-based assay. In our cell-based experiments, at least three biological replicates were analyzed in each single experiment and PBMCs from different donors were used, as indicated in the figure legends. All experiments performed on human tissue samples were conducted in accordance with the Declaration of Helsinki. Human blood was obtained from healthy donors from the Swiss Blood Donation Center in Bern and was used in compliance with the Federal Office of Public Health (authorization no. P_149). The skin was obtained from healthy patients who underwent cosmetic surgery procedures or patients with ACD, or positive patch test reactions to standard contact allergens. The study on human patient samples was approved by the Medical Ethics Committee of the Canton of Bern, Switzerland (no. 088/13; 2019-01068; 2019-00803). Written informed consent was obtained from all patients. Mechanistic studies on blood and human tissue cells were performed using in vitro assays without blinding or randomization.

### Isolation and purification of human T cell subsets from peripheral blood

Peripheral Blood Mononuclear Cells (PBMCs) were isolated according to the Standard Operating Procedure (SOP): PBMC Isolation using SepMate™. Human naïve T cells were isolated from PBMCs using the EasySep™ Human naïve CD4^+^ T Cell Isolation Kit Human (Stemcell Technologies) as per the manufacturer’s instructions. CD4^+^ T cells were isolated from PBMCs using the EasySep positive selection kit (Stemcell Technologies) as per the manufacturer’s instructions. Positively selected CD4^+^ T cells were stained for the subsequent sorting of the T_H_ cell subset. Memory T_H_ cell subsets were sorted with a purity of > 90% according to the expression of chemokine receptors from CD45RA^-^CD25^-^CD8^-^CD3^+^ cells: T_H_1 (CXCR3^+^CCR8^-^CCR6^-^CCR4^-^), T_H_2 (CXCR3^-^CCR8^-^CCR6^-^CCR4^+^), T_H_9 (CXCR3^-^CCR8^+^CCR6^-^CCR4^+^), and T_H_17 (CXCR3^-^CCR8^-^ CCR6^+^CCR4^+^).

### T cell cloning

Individual memory T_H_ cells were directly sorted from CD4^+^ T cells into 96-well plates according to the expression of chemokine receptors (see Isolation and purification of human T cell subsets from peripheral blood). Individual cells were grown by periodic activation with phytohemagglutinin (1 *μ*g/ml; Sigma), and irradiated allogeneic feeder cells (5×10^4^ per well) in a culture medium. Half of the nutrient medium for T cell culture was replaced with a fresh medium every second day, starting from day 2 after reactivation. T_H_ cell clones were analyzed in the resting state (≥ 14 days after the last expansion) or at different time points after polyclonal activation (see T cell culture and activation).

### In vitro T cell differentiation

Naïve T cells were stimulated with *α*CD3/CD2/CD28 beads (T cell/bead = 2:1, Miltenyi) and primed into effector CD4^+^ T cell subsets with no addition of cytokines for T_H_0 cells, IL-12 (5 ng/ml) (BioLegend) for T_H_1 cells, IL-4 (50 ng/ml) (BioLegend) for T_H_2 cells, IL-4 (50 ng/ml) and TGF-*β* (5 ng/ml) (R&D Systems) for T_H_9 cells, and TGF-*β* (5 ng/ml) for iT_REG_. From cell culture initiation to analysis, the culture medium was supplemented with the indicated cytokines every other day. Cells were harvested for RNA sequencing (RNA-seq), quantitative reverse transcription-polymerase chain reaction (RT-qPCR) analysis or intracellular cytokine analysis by flow cytometry at different time points (see below).

### T cell culture and activation

The culture medium consisted of RPMI 1640 with Hepes (Gibco) supplemented with 5% heat-inactivated human serum (Swiss Red Cross, Basel, Switzerland), 1% Glutamax (Gibco), penicillin (50 U/ml) and streptomycin (50 *μ*g/ml) (BioConcept), and IL-2 (50/250 IU/ml) (Roche). The glucose medium consisted of glucose-free RPMI 1640 (Gibco) supplemented with 5% heat-inactivated dialyzed FBS (Gibco), 1% Glutamax (Gibco), penicillin (50 U/ml) and streptomycin (50 *μ*g/ml) (BioConcept), IL-2 (50 IU/ml) (Roche) and glucose (Sigma). T cells were cultured in a 96-well plate with a density of 0.25 × 10^5^ to 1 × 10^5^ cells per well in a total volume of 200 *μ*l of cell culture medium. T cells were polyclonally activated using ImmunoCult Human CD3/CD2/CD28 T Cell Activator (1:100) (Stemcell Technologies).

### Isolation of human T cells from skin biopsies

Lesional and non-lesional skin biopsies of positive patch test reactions to different allergens were cultured in a culture medium as described above, and respective treatments were added to the culture. T cells were harvested and analysed by flow cytometry and RNA-seq.

### Analysis of cytokine expression and S6 phosphorylation by flow cytometry

All antibodies used for flow cytometry are listed in Supplementary Table S1. To analyze cytokine production and S6 phosphorylation, T cells were polyclonally activated using ImmunoCult Human CD3/CD2/CD28 T Cell Activator (1:100) (Stemcell Technologies). Before activation and at different time points thereafter, T cells were additionally stimulated with PMA (50 ng/ml) (Sigma-Aldrich), ionomycin (1 *μ*M) (Sigma-Aldrich), and brefeldin A (10 *μ*g/ml) (Sigma-Aldrich) for 4 h. After viability and surface staining, the cells were fixed and permeabilized using Cytofix/Cytoperm kit (BD Biosciences) as per the manufacturer’s instructions. Fluorescence-labeled antibodies were used to detect intracellular proteins, as well as phosphorylation.

### Proliferation assays

Proliferation assays were performed with carboxyfluorescein diacetate succinimidyl ester (CFSE) staining. The cells were labeled with CFSE (2 *μ*M) in PBS and incubated at 37°C for 8 min. Staining was quenched by adding 2 times the initial staining volume of the cell culture medium and incubating the cells at 37°C for 5 min. After 3 to 4 days, the stained cells were acquired on CytoFLEX (Beckman Coulter), and the data were analyzed using CytExpert (Beckman Coulter) or FlowJo v.10 software (BD Life Sciences).

### Western Blotting

For the analysis of S6 phosphorylation, cells were harvested after treatment, washed with PBS containing the Halt^™^ Protease and Phosphatase Inhibitor cocktail (Thermo Scientific), and lysed in 20 mM Tris-HCl pH 7.5, 0.5% NP40, 25 mM NaCl, and 2.5 mM EDTA containing the Halt^™^ Protease & Phosphatase Inhibitor cocktail (Thermo Scientific). Protein concentration was measured using the Pierce BCA protein assay kit (Thermo Scientific). Samples (5-10 µg of protein per lane) were loaded onto 10% SDS-PAGE gel. For the Western blot analysis of MCT1, 1 × 10^6^ cells were lysed in 30 µl of a sample loading buffer (SLB) consisting of 62.5 mM Tris-HCl (pH 6.8), 2% 2-mercaptoethanol, 2% SDS, 0.02% bromophenol blue, 14.8% Glycerol, and 6M Urea. For the Western blot analysis of PPAR-γ, protein extracts were prepared as follows. Cells were resuspended in an ice-cold lysis buffer (20 mM Hepes, pH 7.8, 10 mM KCl, 1 mM EDTA, 0.65% Nonidet P-40, 1 mM DTT, 1 mM PMSF) and incubated for 15 min on ice. Nuclei were pelleted at 20,000 x g for 20 min at 4°C and lysed in 30 *μ*l SLB. After electrophoresis (150 V, 45 min), proteins were transferred to a 0.45 µm Nitrocellulose Blotting membrane (AmershamTM ProtranTM) by wet transfer (100 V, 75 min). Non-specific sites were blocked for 1 h with 5% non-fat milk in a TBS-T buffer (25 mM Tris, pH 7.5, 150 mM NaCl, and 0.1% Tween 20). Primary antibodies were incubated overnight at 4°C. Membranes were washed with the TBS-T buffer and incubated for 1 h at room temperature with the corresponding secondary antibodies. Using Western Bright Quantum (Advansta) or SuperSignal™ West Atto Ultimate Sensitivity Substrate (Thermo Scientific) the binding of specific antibodies was visualized thereafter using Fusion Pulse TS of Vilber Lourmat (Witec). All antibodies used for Western blotting are listed in Supplementary Table S1.

### RNA sequencing (RNA-seq)

Total RNA was isolated from T cells using the RNeasy kit (Qiagen) as per the manufacturer’s instructions. The samples were submitted to the Next Generation Sequencing (NGS) Platform (Institute of Genetics, University of Bern). RNA integrity was analyzed by Qubit^™^. For all samples, the RNA Integrity Number (RIN) values were ≥ 8. The total RNA was transformed into a library of template molecules using TruSeq® Stranded mRNA Sample Preparation Kits (Illumina®) and the EpMotion 5075 (Eppendorf) robotic pipette system. Single-end 100 bp and paired-end 50 bp sequencing were performed using HiSeq3000 (Illumina®). The RNA-seq reads were mapped to the reference human genome (GRCh38, build 81) using HISAT2 v. 2.0.4 (*39*). HTseq-count v. 0.6.1 (*40*) was used to count the number of reads per gene, and DESeq2 v.1.4.5 (*41*) was used to test for differential expression between groups of samples. The RNA-seq data are deposited on ArrayExpress (accession no. XXX).

### Seahorse

The oxygen consumption rate (OCR) and extracellular acidification rate (ECAR) were measured 24 h after treatment using the Seahorse XFe96 Analyzer (Seahorse Biosciences, Agilent Technologies) as per the manufacturer’s instructions. On the day of the assay, the culture medium was replaced with the Seahorse XF base medium (catalog no. 102353-100), supplemented with reagents necessary to meet the cell culture conditions. Cells were seeded into Seahorse XFe96-well plates (Seahorse Biosciences, Agilent Technologies) with a density of 1.5 × 10^5^ cells per well and a total volume of 50 μl of cell culture medium to obtain eight replicates per condition. Then, cells were equilibrated for 1 h in a non-CO_2_ incubator at 37°C. After measuring the baseline, successive injections of oligomycin (1 μM), carbonyl cyanide-p-trifluoromethoxy-phenylhydrazone (FCCP) (1 μM), rotenone (1 μM), antimycin A (1 μM), and 2-deoxy-d-glucose (2-DG) (50 mM) were delivered to measure the mitochondrial OCR and ECAR. The data were normalized to the DNA content which was determined after each Seahorse assay using CyQUANT™ Cell Proliferation Assay (Thermo Fisher Scientific) as per the manufacturer’s instructions.

### Lactate measurements

T_H_9 clones were pre-incubated with DMSO or BAY-8002 (75 *μ*M) for 1 h before adding IL-9 (5 ng/ml) or culture medium only. After 48 h, the lactate levels were measured using the Lactate-Glo™ Assay (Promega) as per the manufacturer’s instructions. Luminescence was measured using Tecan Reader Spark 10M.

### Glucose measurements

Lesional skin biopsies from positive patch test reactions to different allergens (Supplementary Table S3) and adjacent non-lesional skin biopsies were weighed and resuspended in PBS. The volume of PBS corresponded to 8 times the weight of the biopsy. Skin biopsies were centrifuged at 300 g for 8 min to isolate interstitial fluid of biopsy samples. Glucose concentration in the interstitial fluid was measured using the Glucose-Glo™ Assay (Promega) as per the manufacturer’s instructions. Luminescence was measured using Tecan Reader Spark 10M.

### Glucose Uptake

2-NBDG was used as a tool to study cellular glucose uptake. Cells were washed in PBS and incubated with 1 ng/ml 2-NBDG (Cayman) in a glucose-free medium for 15 min at 37°C in 5% CO_2_. After washing, cells were either analyzed by flow cytometry or sorted into three different populations (low, medium and high) according to their 2-NBDG uptake rate. RNA was isolated for the RT-qPCR analysis.

### Immunofluorescence

Skin biopsies of ACD patients were embedded in paraffin, cut into 4 *μ*m thick sections, and heated for 20 min at 63°C. The samples were stained using a BOND Autostainer and included dewaxing, pre-treatment with a buffer pH 9 for 20 min at 95°C, and sequential double staining. Skin biopsies from positive patch test reactions were placed in an optimal cutting temperature (OCT) compound, snap-frozen, and stored at -80°C. Samples were cut into 6 *μ*m cryo-sections, fixed with acetone at 4°C for 10 min, and blocked with normal goat serum (1:50) (Dako) at room temperature for 15 min. Primary antibodies were added at room temperature for 60 min, followed by washing with TBS-Saponin 0.1%. Secondary antibodies were also added at room temperature for 60 min, followed by washing with TBS-Saponin 0.1%. Slides were mounted using Fluoromount-G™ Mounting Medium with DAPI (Southern Biotech).

### Quantitative reverse transcription-polymerase chain reaction (RT-qPCR)

Total RNA was isolated from cultured in vitro or in vivo primed T cells or ex vivo sorted T cells using the RNeasy kit (Qiagen) as per the manufacturer’s instructions. RNA from snap-frozen skin biopsies was isolated using the RNeasy Lipid Tissue Kit (Qiagen) as per the manufacturer’s instructions. The total mRNA quality was measured using the ND-1000 Spectrophotometer (Thermo Fisher Scientific) or 2100 Bioanalyzer (Agilent). Complementary DNA was generated using Omniscript reverse transcriptase (Qiagen). Real-time PCR was performed using TaqMan probe-based assays and measured using the 7300 Real-Time PCR System (Applied Biosystems). The expression of each ligand transcript was determined relative to the reference gene transcript (HPRT-1) and normalized to the expression of the target gene using the 2^−ΔΔCt^ method. The data are represented as arbitrary relative units. All used primers were acquired from Thermo Fisher and are listed in Supplementary Table S2.

### Statistical analysis

Statistical analysis was performed using the GraphPad Prism 9 software. For between-group comparisons, a one-way or two-way analysis of variance (ANOVA) was used, followed by pairwise comparisons for each group and either a Dunnett’s, Tukey’s or Šidák’s test to correct for multiple testing. The matched samples were analyzed using two-tailed paired t-tests or repeated-measures ANOVA whereas unmatched samples were analyzed with two-tailed unpaired t-test. The n values and the respective statistical methods for individual experiments are indicated in the figure legends. For all statistical analyses, a 95% confidence interval and p-value < 0.05 were considered significant.

## Abbreviations

ACD: Allergic contact dermatitis
CFSE: Carboxyfluorescein succinimidyl ester
ECAR: Extracellular acidification rate
EoE: Eosinophilic esophagitis
FCCP: Carbonyl cyanide-p-trifluoromethoxyphenylhydrazone
Glc: Glucose
HIF-1α: Hypoxia-inducible factor 1-alpha
IL: Interleukin
ILC2: Type 2 innate lymphoid cell
MCT1: Monocarboxylate transporter 1
NL: Non-lesional
OCR: Oxygen consumption rate
PBMCs: Peripheral Blood Mononuclear Cells
PMA: Phorbol 12-myristate 13-acetate
PPAR-γ: Peroxisome proliferator-activated receptor-γ
PPREs: PPAR response elements
RT-qPCR: Quantitative reverse transcription-polymerase chain reaction
RXR: Retinoic X receptor
TCR: T cell receptor
T_EM_: Effector memory T helper cells
T_H_: T helper
pT_H_2: Pathogenic T_H_2 cells
mTORC1: Mammalian target of rapamycin complex 1
2-DG: 2-Desoxy-D-glucose

## Supplementary Materials

### Supplementary Methods

RNA-seq of T_H_ cell subsets primed in vitro

RNA-seq of non-lesional and lesional skin biopsies post allergen application

RNA-seq of T_H_9 clones in presence of the PPAR-γ inhibitor GW9662

RNA-seq of IL-9R^+^ T_H_ cells in presence of recombinant human IL-9

### Supplementary Figures

Fig. S1: In vitro and in vivo primed T_H_9 cells show the key features of pathogenic T_H_2 cells

Fig. S2: PPAR-γ mediates the high glycolytic activity of T_H_9 cells

Fig. S4: mTORC1 integrates bioenergetics with the effector function in T_H_9 cells

### Supplementary Tables

Table S1: Antibodies, recombinant proteins and chemicals used in this study

Table S2: Quantitative reverse transcription-polymerase chain reaction (RT-qPCR) primers used in the study

Table S3: Allergens from Figure 6D

## Acknowledgements

We thank I.K. and the genomics core facility of the Department of Clinical Research, University of Bern for technical support. We are grateful to Ben Roediger and Curdin Conrad for their helpful comments.

## Author contributions

O.S., N.L.B., F.L., N.B., L.v.M., C.B., M.L., and C.Sc. designed and performed the experiments; O.F., S.F., A.F., JM. N., M.L., and C.Sc. provided essential reagents and funding; M.P.G, C.L., M.B., D.S., and C.Sc. provided patient samples; O.S., N.L.B., F.L., L.v.M., and C.Sc analyzed the data; F.L., N.L.B., and C.Sc. wrote, edited, and revised the article.

## Data and materials availability

The RNA-seq data are deposited on ArrayExpress (accession no. E-MTAB-5739). N.L.B., F.L., C.B., and C.Sc are members of the SKINTEGRITY.CH collaborative research project.

