## Supplementary Material for "PPAR-γ regulates the effector function of human T_H_9 cells by promoting glycolysis"

### Supplementary Methods

**RNA-seq of T_H_ cell subsets primed in vitro**

Naïve T-cells were isolated from PBMCs by EasySep negative selection kit (Stemcell Technologies) according to the manufacturer’s instructions. Naive T_H_ cells were differentiated under T_H_1 (IL-12), T_H_2 (IL-4), T_H_9 (IL-4+TGF-β), or iT_REG_ (TGF-β) priming conditions for 7 days (see Materials and Methods, In vitro T cell differentiation). Total RNA was isolated from clones with the RNeasy Micro Kit (Qiagen) according to manufacturer’s instruction and RNA-seq was performed.

**RNA-seq of non-lesional and lesional skin biopsies post allergen application**

Six donor-matched skin biopsies were taken from non-lesional (NL) and lesional skin of positive patch-test reactions to nickel at 24h, 48h, and 120h post allergen application, respectively. Skin biopsies from positive patch test reactions were placed in an optimal cutting temperature (OCT) compound, snap-frozen, and stored at -80°C. Total RNA was isolated from biopsies with the RNeasy Lipid Tissue Mini Kit (Qiagen) according to manufacturer’s instruction and RNA-seq was performed.

**RNA-seq of T_H_9 clones in presence of the PPAR-γ inhibitor GW9662**

Human CD4^+^ T cells were isolated from PBMCs by EasySep positive selection kit (Stemcell Technologies) according to the manufacturer’s instructions. Positively selected CD4^+^ T cells were stained for subsequent T_H_ cell subset sorting (see Materials and Methods, Isolation and purification of human T cell subsets from peripheral blood). Total RNA was isolated from T_H_9 clones after incubation with DMSO or GW9662 for 48h followed by 12h activation with anti-CD3/CD2/CD28 with RNeasy Micro Kit (Qiagen) according to manufacturer’s instruction.

**RNA-seq of IL-9R^+^ T_H_ cells in presence of recombinant human IL-9**

Human CD4^+^ T cells were isolated from PBMCs by EasySep positive selection kit (Stemcell Technologies) according to the manufacturer’s instructions. Positively selected CD4^+^ T cells were stained for subsequent T_H_ cell subset sorting (see Materials and Methods, Isolation and purification of human T cell subsets from peripheral blood). After T cell single cell cloning, cells were screened for IL-9R expression by flow cytometry. Skin biopsies from positive patch test reactions were cultured in culture medium and CD4^+^ T cells were isolated and screened for IL-9R expression by flow cytometry. Five IL-9R^+^ T_H_ clones isolated from blood and three IL-9R^+^ T_H_ cells isolated from skin biopsies, respectively, were incubated in presence or absence of recombinant human IL-9 (5ng/ml) for 12h. Total RNA was isolated with the RNeasy Micro Kit (Qiagen) according to manufacturer’s instruction and RNA-seq was performed.

### Supplementary Figures

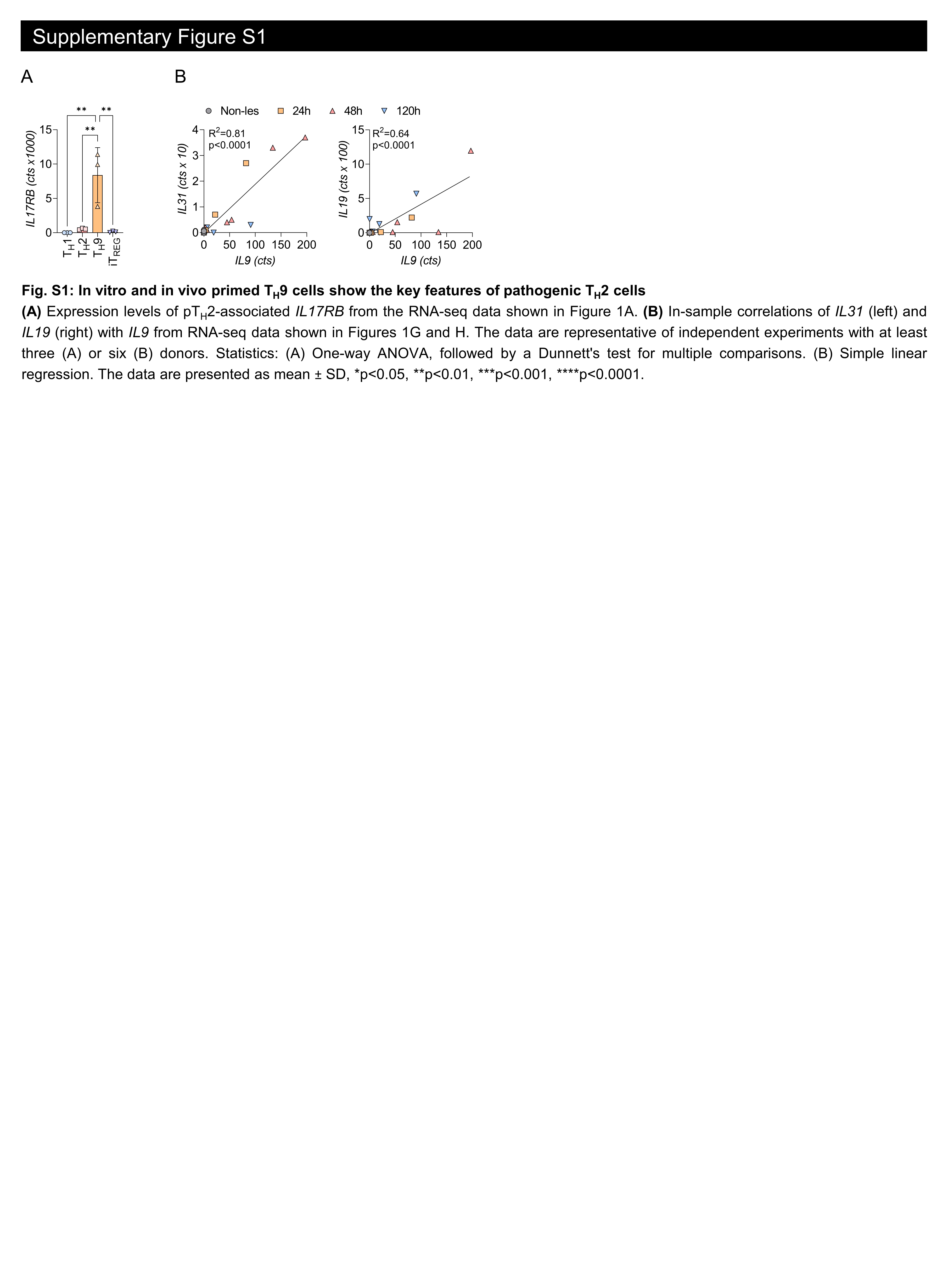

**Fig. S1: In vitro and in vivo primed T_H_9 cells show the key features of pathogenic T_H_2 cells**

**(A)** Expression levels of pT_H_2-associated *IL17RB* from the RNA-seq data shown in Figure 1A. **(B)** In-sample correlations of *IL31* (left) and *IL19* (right) with *IL9* from RNA-seq data shown in Figures 1G and H. The data are representative of independent experiments with at least three (A) or six (B) donors. Statistics: (A) One-way ANOVA, followed by a Dunnett's test for multiple comparisons. (B) Simple linear regression. The data are presented as mean ± SD, *p<0.05, **p<0.01, ***p<0.001, ****p<0.0001.

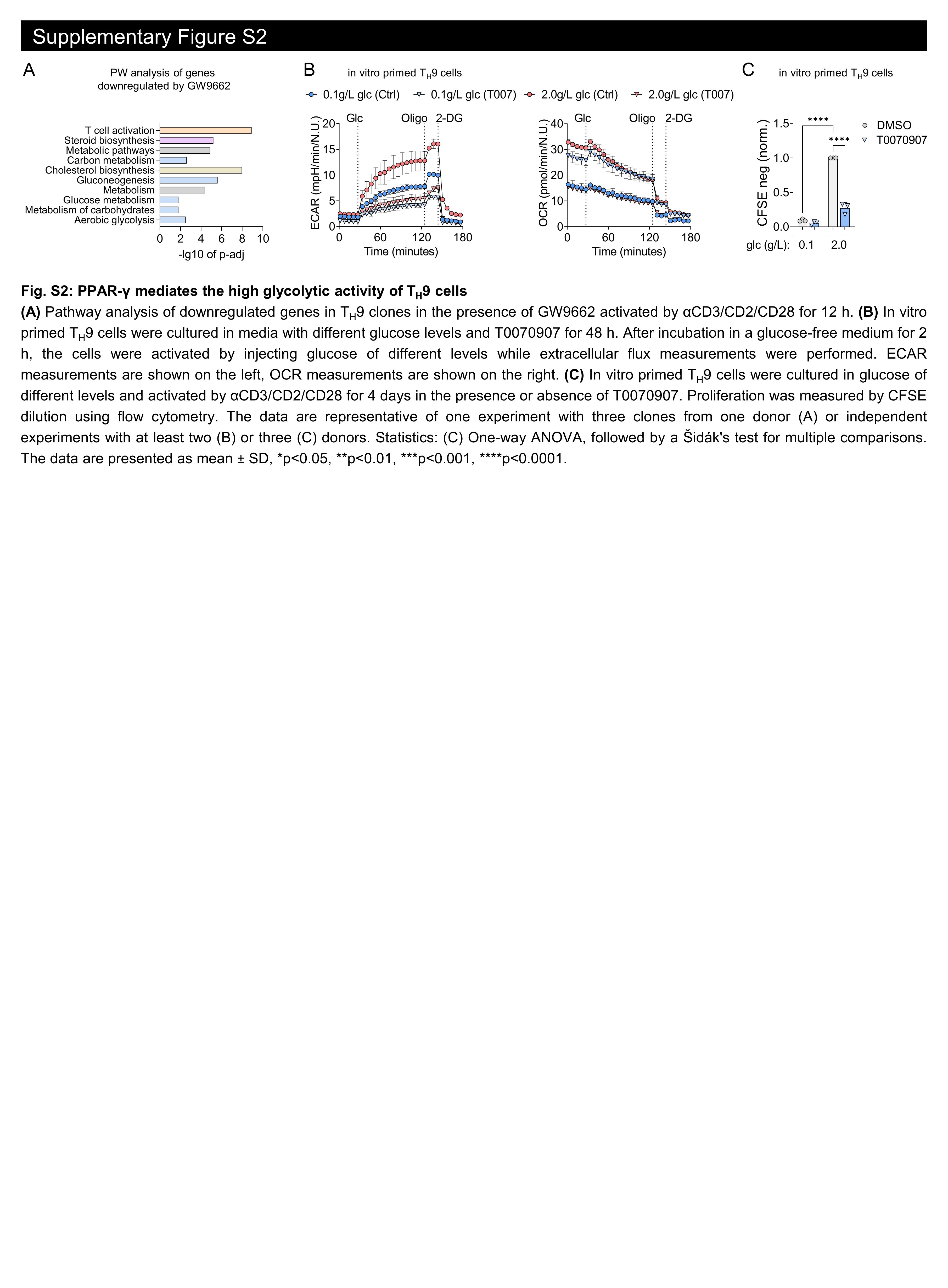

**Fig. S2: PPAR-γ mediates the high glycolytic activity of T_H_9 cells**

**(A)** Pathway analysis of downregulated genes in T_H_9 clones in the presence of GW9662 activated by αCD3/CD2/CD28 for 12 h. **(B)** In vitro primed T_H_9 cells were cultured in media with different glucose levels and T0070907 for 48 h. After incubation in a glucose-free medium for 2 h, the cells were activated by injecting glucose of different levels while extracellular flux measurements were performed. ECAR measurements are shown on the left, OCR measurements are shown on the right. **(C)** In vitro primed T_H_9 cells were cultured in glucose of different levels and activated by αCD3/CD2/CD28 for 4 days in the presence or absence of T0070907. Proliferation was measured by CFSE dilution using flow cytometry. The data are representative of one experiment with three clones from one donor (A) or independent experiments with at least two (B) or three (C) donors. Statistics: (C) One-way ANOVA, followed by a Šidák's test for multiple comparisons. The data are presented as mean ± SD, *p<0.05, **p<0.01, ***p<0.001, ****p<0.0001.

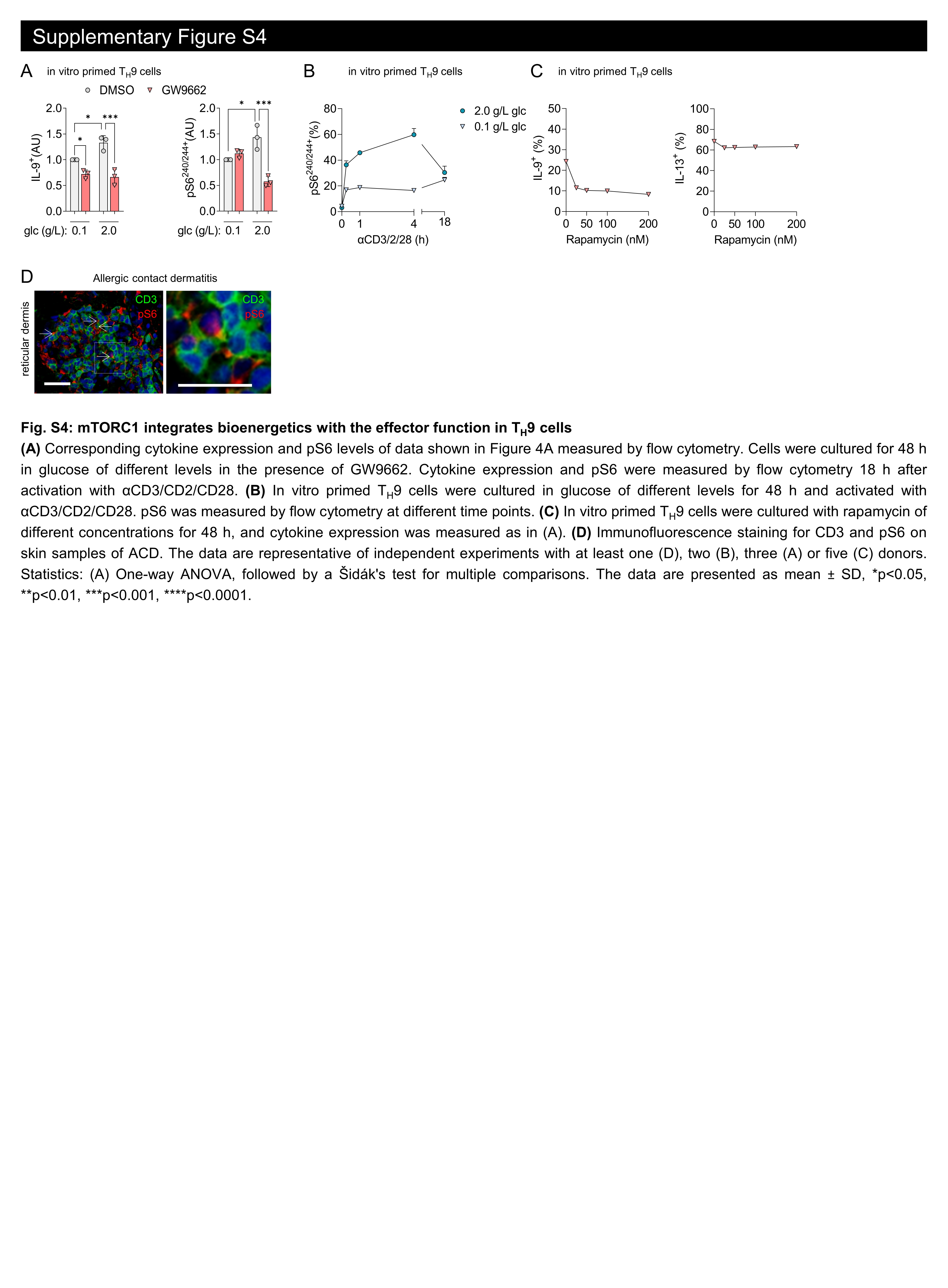

**Fig. S4: mTORC1 integrates bioenergetics with the effector function in T_H_9 cells**

**(A)** Corresponding cytokine expression and pS6 levels of data shown in Figure 4A measured by flow cytometry. Cells were cultured for 48 h in glucose of different levels in the presence of GW9662. Cytokine expression and pS6 were measured by flow cytometry 18 h after activation with αCD3/CD2/CD28. **(B)** In vitro primed T_H_9 cells were cultured in glucose of different levels for 48 h and activated with αCD3/CD2/CD28. pS6 was measured by flow cytometry at different time points. **(C)** In vitro primed T_H_9 cells were cultured with rapamycin of different concentrations for 48 h, and cytokine expression was measured as in (A). **(D)** Immunofluorescence staining for CD3 and pS6 on skin samples of ACD. The data are representative of independent experiments with at least one (D), two (B), three (A) or five (C) donors. Statistics: (A) One-way ANOVA, followed by a Šidák's test for multiple comparisons. The data are presented as mean ± SD, *p<0.05, **p<0.01, ***p<0.001, ****p<0.0001.

**Supplementary Tables**

Table S1: Antibodies, recombinant proteins and chemicals used in this study

| Flow cytometry - Surface staining | Antibody | Clone | Conjugation | Company | Dilution |
| --- | --- | --- | --- | --- | --- |
|  | Mouse anti-human CXCR3, mAb | G025H7 | AF647  FITC | biolegend (353712)  biolegend (353704) | 1:60  1:200 |
|  | Mouse anti-human CD45RA, mAb | HI100 | APC-Cy7  PerCP-Cy5.5 | biolegend (304128)  biolegend (304122) | 1:200  1:1600 |
|  | Mouse anti-human CD8, mAb | RPA-T8 | FITC  PerCP-Cy5.5  BV660 | biolegend (301021)  biolegend (344710)  biolegend (301042) | 1:100  1:1600  1:400 |
|  | Mouse anti-human CD25, mAb | BC96 | FITC | biolegend (302604) | 1:100 |
|  | Mouse anti-human CCR8, mAb | L263G8 | PE  APC | biolegend (360604)  biolegend (360609) | 1:60  1:200 |
|  | Mouse anti-human CCR7, mAb | G043H7 | PE-Cy7 | biolegend (353226) | 1:200 |
|  | Mouse anti-human CCR4, mAb | L291H4 | PE-Cy7  BV605 | biolegend (359410)  biolegend (359417) | 1:60  1:200 |
|  | Mouse anti-human CCR6, mAb | G034E3 | PerCP-Cy5.5  BV421 | biolegend (353406)  biolegend (353407) | 1:60  1:200 |
|  | Mouse anti-human CD3, mAb | OKT3 | BV785 | biolegend (317330) | 1:200 |
|  | Mouse anti-human CD4, mAb | OKT4 | APC-Cy7 | biolegend (317418) | 1:400 |
|  | Mouse anti-human IL-9R, mAb | AH9R7 | PE | biolegend (310404) | 1:200 |
|  | Mouse anti-human IgG2b, mAb | MG2b-57 | PE | biolegend (401207) | 1:200 |
|  | 2-NBDG |  | FITC | Cayman Chemicals | 1ng/ml |
|  | CFSE |  | FITC | Selleckchem | 2uM |
| Flow cytometry – Intracellular staining | Antibody | Clone | Conjugation | Company | Dilution |
|  | Rat anti-human IL-4, mAb | MP4-25D2 | PE-Cy7 | biolegend (500824) | 1:400 |
|  | Rat anti-human IL-5, mAb | TRFK5 | BV421 | biolegend (504311) | 1:400 |
|  | Mouse anti-human IL-9, mAb | MH9A4 | PE | biolegend (507605) | 1:400 |
|  | Rat anti-human IL-13, mAb | JES10-5A2 | APC | Biolegend (501907) | 1:300 |
|  | Mouse anti-human INF-γ, mAb | B27 | PE-Cy7 | Biolegend (506518) | 1:400 |
|  | Rabbit anti-human pS6, mAb | D57.2.2E | AF488 | Cell Signaling (4803S) | 1:800 |
| Immunofluorescence Assays | Antibody | Clone | Conjugation | Company | Dilution |
|  | Mouse anti-human CD3, mAb | F7.2.38 | unconjugated | Dako (M7254) | 1:50 |
|  | Mouse anti-human CD4, mAb | 4B12 | unconjugated | Novocastra (NCL-L-CD4-368) | 1:50 |
|  | Mouse anti-human PPAR-γ, mAb | E8 | unconjugated | Santa Cruz (sc-7273) | 1:100 |
|  | Rabbit anti-human pS6, mAb | D57.2.2E | unconjugated | Cell Signaling (4858S) | 1:100 |
|  | Goat anti-rabbit IgG, pAb |  | AF488 | Invitrogen (A11008) | 1:500 |
|  | Goat anti-rabbit IgG, mAb |  | AF594 | Invitrogen (A11072) | 1:500 |
|  | Goat anti-mouse IgG1, mAb |  | AF594 | Invitrogen (A21125) | 1:500 |
|  | Fluoromount G with DAPI |  |  | Southern Biotech (0100-20) |  |
| Western Blotting | Antibody | Clone | Conjugation | Company | Dilution |
|  | Rabbit anti-human PPAR-γ, mAb | C26H12 | unconjugated | Cell Signaling (2435) | 1:500 |
|  | Rabbit anti-human pS6, mAb | D57.2.2E | unconjugated | Cell Signaling (4858S) | 1:2000 |
|  | Rabbit anti-human S6, mAb | 5G10 | unconjugated | Cell Signaling (2217S) | 1:1000 |
|  | Mouse anti-human MCT1, mAb | H-1 | unconjugated | Santa Cruz (sc-365501) | 1:250 |
|  | Rabbit anti-human Histone H2B, pAb |  | unconjugated | Sigma (SAB4502231) | 1:1000 |
|  | Mouse anti-human F-Actin, mAb | ACTN05 | unconjugated | Invitrogen (MA5-11869) | 1:6000 |
|  | Goat anti-mouse IgG, pAb |  | HRP | Thermo Fisher (G21040) | 1:5000 |
|  | Goat anti-rabbit IgG, pAb |  | HRP | Fisher scientific (31462) | 1:5000 |
| Recombinant proteins & Chemicals | Name |  | Format | Company | Final Concentration |
|  | rhIL-2 |  | purified | Hoffmann-La Roche | 50/250 IU/ml |
|  | rhIL-4 |  | purified | biolegend (574006) | 50ng/ml |
|  | rhTGF-beta |  | purified | R&D (240-B-010) | 5ng/ml |
|  | rhIL-12 |  | purified | biolegend (573004) | 5ng/ml |
|  | rhIL-9 |  | purified | biolegend (594404) | 5ng/ml |
|  | GW9662 |  |  | Sigma-Aldrich (M6191) | 10uM |
|  | T0070907 |  |  | Selleckem (S2871) | 10uM |
|  | Rapamycin |  |  | Sigma-Aldrich (R0395) | 50nM |
|  | BAY-8002 |  |  | Selleckem (S8747) | 75uM |
|  | 2-DG |  |  | Sigma-Aldrich (D8375) | 1mM |

Table S2: Quantitative reverse transcription-polymerase chain reaction (RT-qPCR) primers used in the study

| Gene Transcript | Gene | Species | TaqMan Primer |
| --- | --- | --- | --- |
| IL-4 | *IL4* | human | Hs00174122_m1 |
| IL-5 | *IL5* | human | Hs01548712_g1 |
| IL-9 | *IL9* | human | Hs00174125_m1 |
| IL-13 | *IL13* | human | Hs00174379_m1 |
| PPAR-gamma | *PPARG* | human | Hs01115513_m1 |
| HPRT1 | *HPRT1* | human | Hs99999909_m1 |

Table S3: Allergens from Figure 6D

| Sample ID | Allergen | Intensity of Reaction |
| --- | --- | --- |
| 1 | Perubalsam | +++ |
| 2 | Perubalsam | +++ |
| 3 | p-Phenylendiamin | +++ |
| 4 | Primin | +++ |
| 5 | Propolis | +++ |
| 6 | p-Phenylendiamin | +++ |
| 7 | p-Phenylendiamin | +++ |
